# The Roboscope: Smart and Fast Microscopy for Generic Event-Driven Acquisition

**DOI:** 10.1101/2024.09.24.614735

**Authors:** Julia Bonnet, Youssef El-Habouz, Célia Martin, Maelle Guillout, Louis Ruel, Baptiste Giroux, Claire Demeautis, Benjamin Mercat, Otmane Bouchareb, Jacques Pécreaux, Marc Tramier

## Abstract

Automation of fluorescence microscopy is a challenge for capturing rare or transient events in biology and medicine. It relies on smart devices that integrate and interpret the observed data, and react to the targeted biological event. We report on the Roboscope, a novel autonomous microscope combining sequence interruption and deep learning integration, allowing generic event-driven acquisitions. This system distinguishes itself by its adaptability to various experiments, quick capture of dynamic events, and minimal data greediness – training with less than 100 images per class. The Roboscope’s capability is demonstrated in non-synchronized cells by capturing the metaphase, a 20-minute event happening once per day or less. Conversely, double thymidine-block synchronisation, despite occurring during DNA replication, may perturb mitotic-spindle mechanics. The Roboscope’s versatility and efficiency offer significant advancements to tackle the current challenges of cell biology, spreading out advanced microscopy methods to fundamental research as well as high content screening and precision medicine.

## Introduction

Fluorescence microscopy, through its diverse modalities, offers an unparalleled approach to living. However, despite the microscopes being fully motorised, complex experiments still require expert supervision; acquisitions of rare events or high content screening are achieved by acquiring continuously or at multiple time points, illuminating the sample uselessly and generating a flood of unqualified data. Alternatively, an experimenter needs constant monitoring over the extended duration of experiments to qualify the events of interest (EOI). This selection by a scientist who knows the tested hypothesis may introduce a bias. The automation of fluorescence microscopy is thus strategic to increase robustness and numbers, and boost the impact of cellular imaging in biological research. The pioneering work of Micropilot paved the way for image-processing-aided acquisition, alternating phases of sample screening for EOIs and image processing, and then the specific acquisition of EOIs if applicable^1^. Beyond finding EOIs, early days setups, dedicated to a specific experiment, also spared the sample from excessive illumination and thus photodamages^2^. In recent years, taking advantage of the widespread use of Artificial Intelligence (AI) for data analysis in biology^3-7^, frameworks were proposed to perform real-time grabbed-image processing leaving to the user the implementation of the feed-back to the microscope^8-10^. Meanwhile, several groups have developed smart microscopes for event-driven acquisition dedicated to specific modalities, like wide-field^11^, stimulated emission depletion^12^, structured illumination microscopy^13^, super-resolution localisation microscopy^14^, or lattice light sheet^15^. These solutions share the same limitation; they are heavily tailored to their attached microscopy setup and biological application. Our work aimed to increase the user-friendliness and the genericity of smart microscopes.

We present our solution, the Roboscope. It combines automation with innovative imaging-sequence interruption and task parallelisation to ensure the real-time acquisition of EOI. A novel AI algorithm ensures the genericity of EOI detection, keeping the training set small. We used embedded technology based on a modular system provided by Inscoper (I.S. Imaging Solution)^16^ to ensure fast driving on the microscope. Its operation involves a patented sequence interruption approach that breaks a first acquisition screening for EOIs to immediately perform specific imaging with a distinct microscope configuration^17^.

As proof of concept for this novel system, we set out to investigate how synchronising cells could impact the mechanics of the metaphasic spindle. We used the spindle length as a proxy for the dynamics of spindle components^18-22^. We restricted our investigation to double thymidine block synchronisation here, as it acts long before mitosis^23^. To investigate non-synchronised cells, we instructed the Roboscope to detect metaphase in a field of living cells using nucleus labelling, interrupt this screening, modify the optical acquisition parameters to focus on a single cell and acquire fluorescently labelled spindle poles in a different wavelength at high frame rate. Once done, the Roboscope continues screening to find other metaphasic cells. Our generic and user-friendly approach will enable repeating such EOI detection in multiple contexts.

## Results

### Hardware architecture of the smart microscope to achieve task parallelisation

To achieve generic and real-time EOI detection and acquisition, we assembled task-dedicated modules (Fig 1) to get our smart system, contrasting with the current design featuring an all-purpose computer connected to a microscope and a scientific camera. Key to success, it featured a dedicated device running a generic AI image-analysis algorithm. To optimise the execution speed, we executed demanding tasks in parallel on a dedicated microcontroller. It, therefore, implied tight management of interruptions. To do so and efficiently drive the microscope, we built on the solution sold by the Inscoper company (I.S. Imaging Solution, Inscoper). Of notice, this latter system dissociates the device control from the user interface; it uses a dedicated embedded system to manage the device independently from the user interface, image capture, and storage, tasked to the computer^16,24^. This specialised embedded system works autonomously during the acquisition process using parallel and bi-directional communication with the different microscope devices for optimal speed. Furthermore, its broad device library ensures one can control most camera-based microscopes, allowing the Roboscope to be adapted to a vast range of setups. Based on the same idea, we embedded the image processing on a dedicated device, where a graphics processing unit (GPU) enabled massively parallel computing. The CPU of this device was tasked to drive the camera to grab the images. Meanwhile, the driving of all the microscope devices is carried out by the Inscoper device controller. This architecture ensured optimal performance by distributing the most demanding tasks, namely the device driving and image processing, to the two dedicated processing units, while the computer barely ran the user interface and gathered the qualified images.

**Figure 1:**
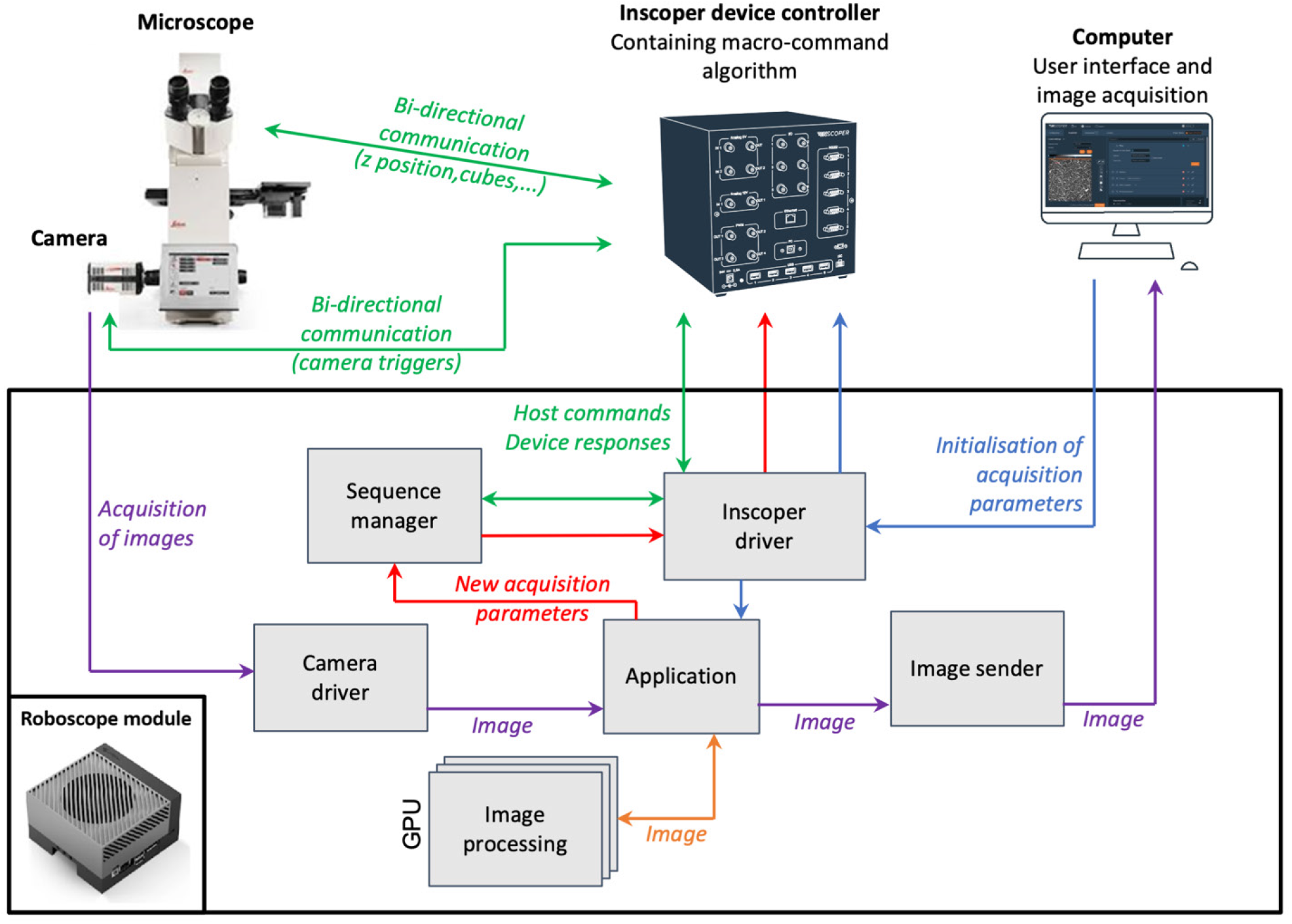
The control system of the smart microscope. The Roboscope setup featured 4 modules: a camera-based microscope, a device controller, a Roboscope module and a computer. The Inscoper Imaging Solution drove the fluorescence microscope through bidirectional communication (green arrows). Before launching the acquisition, the parameters captured from the user interface were sent to the Roboscope module (blue arrows), which centralised communication with the computer, and partly forwarded them to the controller to set up the experiment. The Roboscope module, an Nvidia AGX Xavier GPU embedded system (black box), retrieved the images from the camera (purple arrows depicting image flow), analysed them on the fly (grey shaded boxes detailing the modular architecture of the software) and forward them if required to the computer. Image processing is fostered by artificial intelligence accelerated through GPU computing (Orange arrow). Upon processing results, new acquisition parameters are sent to the controller (red arrows). Within the Roboscope module (grey shaded boxes), the software is assembled by sub-modules to enable parallelised tasks. The *application* sub-module is the organising centre: upon receiving images from the *camera driver*, it submits their analysis to *image processing* and retrieves the probability of each class, then decides through conditionals whether to change the acquisition sequence. If so, it communicates adequately with the *sequence manager*. Several sub-modules are tasked specifically to communicate with devices.

### Software architecture of the Roboscope module

Fast and smart acquisition relies on real-time processing of the images grabbed by the camera. To do so, we designed a dedicated software architecture for the Roboscope module — the system’s brain (Fig 1, black frame). We enabled parallel execution through software modularity: one sub-module per hardware-module connection or per analysis task, coordinated by the *application* sub-module. Users can design their application, including image processing, the targeted event, and its consequential action, without coding. The exact program executed in these modules depends on the application and processing of interest. It is selected from a catalogue and parametrised without coding via the user interface on the computer; next, appropriately compiled code is loaded in the Roboscope module before starting the acquisition; it can then be used to execute event-driven acquisitions. During these, the images are received via the *camera driver* sub-module and sent to the *application* sub-module, which will redistribute them depending on the application chosen to the *image processing* sub-module and/or user interface via the *image sender* sub-module. As soon as an EOI is detected, the running sequence is interrupted; new acquisition parameters are calculated by the *sequence manager* sub-module and sent to the device controller via the *Inscoper driver* sub-module. Such an architecture ensures fast event-driven acquisitions while being amenable to achieving multiple types of experiments.

### A generic and no-code event detection system

To ensure generic and no-code EOI detection, we embedded dedicated and optimised pieces of software in the *image processing* sub-module to be invoked by the *application* sub-module. They can be either implemented via traditional coding or artificial intelligence. In our exemplar application, images of large fields of nucleus-labelled cells were pre-processed for standardisation of intensities, and then individual cell vignettes were obtained using conventional pre-processing by intensity threshold (Suppl Info). Such an approach turned out generic enough for the various cell ranges we tested and computed quickly. Notably, the generic architecture of our solution permits embedding a specific deep learning network for more complex segmentation, like cells in tissue^25^. EOIs from cell vignettes were detected by our semi-supervised generative adversarial network (sGAN) (El Habouz et al., in preparation; see Suppl Info for details). In a nutshell, we combined a generative model, classically featuring a generator and a discriminator network, and a supervised classifier, which shared its weights with the discriminator for all layers but the last one^26-28^ (Fig S1). The discriminator and supervised classifier were only 20-layer depth (Table S1), ensuring fast classification when embedded into the Roboscope and reducing the training set’s size. We trained our sGAN on 400 labelled vignettes (80 per mitosis phase class) and 185 non-labelled ones from the public database mitocheck^29^ by concurrently optimising both the GAN part and the supervised classifier part. We used data augmentation to make our classifier insensitive to changes in position, orientation, or illumination intensity (Suppl Info). We then performed domain adaptation to vignettes produced on Zeiss and Leica microscopes used in our experiment using the same cell line and labelling as in our application (Suppl Info). Training is achieved on a GPU-powered workstation, and the trained network is then embedded in the roboscope module. It ensured a generalisable real-time classification with a small dataset using transfer learning or fine-tuning (El Habouz et al., in preparation; see Suppl Info for details). To ensure real-time classification during the experiments, we benefited from the GPU of the embedded system. It is noteworthy that our architecture is flexible enough to enable distributed computing of the image processing to a dedicated high-performance system through the network if image processing is too demanding. In a broader take, the Roboscope, through its dedicated hardware, modular software architecture, and innovative embedded deep network, ensured fast event-driven acquisition of EOIs, preserving genericity to adapt to a broad range of applications with no code and reduced learning datasets.

### Sequence interruption to detect metaphase, a rare mitotic event

To showcase the functioning of our smart microscopy, we investigated the simple and iconic model of imaging mitotic events during the cell cycle. We set up the acquisition of U2OS cell nuclei labelled with Hoechst 33342 to identify metaphase events. We also transfected the cells with an eGFP-Centrin2 plasmid to monitor the mitotic spindle behaviour and dynamics through its pole-to-pole length^30^. In contrast with the previous autonomous microscopes, we executed the acquisition and the image processing in parallel (Fig 2A). We thus designed an automation technique to interrupt the first sequence (screening the sample for EOI) to perform a second sequence of commands to acquire the detected EOI^17^ (Fig 2B). Importantly, this second sequence used parameters dynamically supplied by the image processing. Multiple EOIs can be handled by several sequences #2 performed sequentially. Because it caused a delay between detection and processing of the n^th^ hit, we thus set an expiration time for each hit, namely 10 minutes, in our example, since metaphase is transient. At the end, sequence #1 is resumed to target other events. In further detail, during sequence #1, we acquired images of cell nuclei at low magnification (20x) and 100 ms camera exposure once per 10 minutes (Fig 2B1). In sequence #2, we performed a fast time-lapse at 10 Hz (every 100 ms with a camera exposure of 70 ms) for typically 6 minutes was acquired at higher magnification (63x) to monitor the centrosomes (Fig 2B2). The transition from sequence #1 to sequence #2 included a centring of the field of view on the identified metaphasic cell using localisation information from image processing, a change of lens from 20x to 63x and a modification of the fluorescence filter set from Hoechst fluorescence to GFP fluorescence. Notably, an autofocus technical sequence was intercalated in between #1 and #2; it featured grabbing 10 planes spaced by 1 µm to determine and set the z-position with the best contrast (sharpness). About the EOI finding, the detection of metaphasic cells was performed on 72×72 pixels grey-scaled single-cell vignettes showing the nucleus by our trained sGAN. It classifies the images into 7 classes, providing the events’ probability and statistics on all phases (Fig 2B3, Suppl Info). If metaphase probability is above a given threshold, the cell is considered as a valid EOI. The pre-trained model described above was domain-adapted by transfer learning with 768 cell vignettes grossly equilibrated per class, data-augmented. It then achieved 79 % accuracy on the test with a set of 280 vignettes (Suppl Info, Fig S7). Finally, our sGAN ran on an Nvidia AGX Xavier and took 10 ± 0.6 ms to classify a single vignette (mean ± SD, n = 13264), while segmentation of the large field took 696 ms. With about 400 cells per image, it summed up to 4.9 s per frame, enabling the detection of EOI faster than repetition time during screening. With this approach, we grab the EOI (metaphasic cell) as fast as possible after its detection, ensuring the acquisition of rare and brief mitotic events. It exemplified the functioning of the Roboscope, although our system is much more versatile.

**Figure 2:**
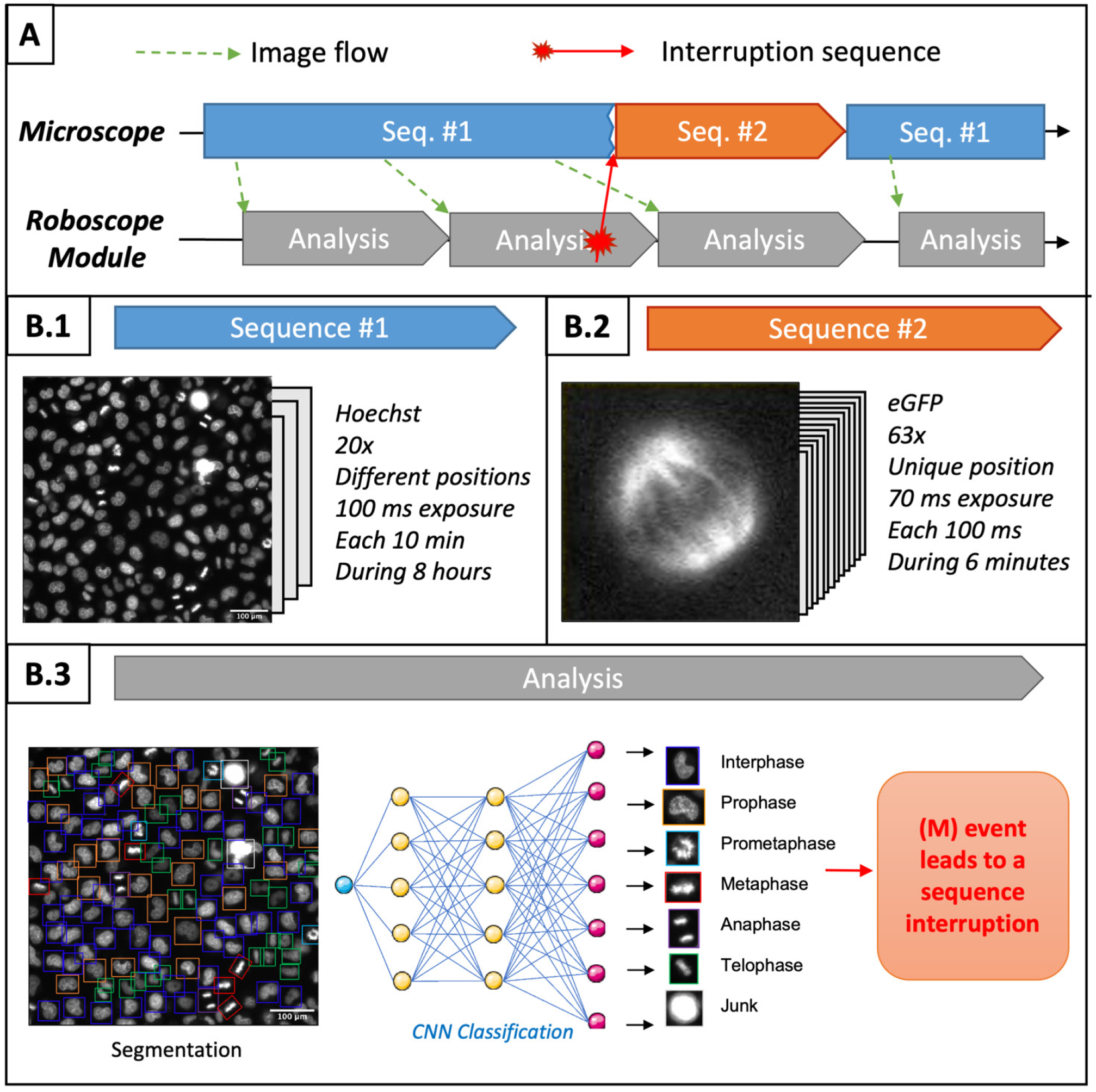
Time diagram detailing events during Roboscope acquisition. **(A)** On the upper timeline, the microscope produces images through sequence 1. (Green arrows) They are sent to the Roboscope module (lower timeline) and analysed concurrently. Once an EOI is detected, (Red arrow) the module instructs the microscope controller to interrupt sequence #1 and start sequence #2. This latter depends on parameters retrieved through the analysis. Upon completion, the roboscope module continues the sequence #1 back. **(B)** Detailed tasks: **(B.1)** Exemplar sequence #1: Cells with Hoechst nuclear staining imaged using a 20x objective and DAPI filter to screen the sample at different positions at one repetition every 10 min. **(B.2)** Exemplar sequence #2: GFP stained microtubule of metaphasic cells displaying the spindle imaged using a 63x objective and FITC filter at 10 frames per second for 6 min. **(B.3)** Typical cell classification by the image processing submodule within the Roboscope unit. In each image, interphase and mitotic cells are segmented and classified with sGAN into 6 stages of cell division. The last class, junk, is also used to detect possible abnormalities.

### Characterising the effect of synchronisation on the mechanics of the metaphasic spindle

Equipped with our event-driven acquisition system, we further investigated mitosis without synchronising cells. Mitosis is classically the last step of the cell cycle and is characterised by consecutive and morphologically distinct stages driven by centrosomes and microtubule organizations^31^. Defects in mitosis, especially those leading to aneuploidy of the daughter cells, contribute to tumour development and resistance to treatments^32,33^; targeting the mechanisms responsible for chromosome segregation is one of the most effective strategies used in cancer chemotherapy^34,35^. Importantly, to understand the underlying mechanisms, the dynamics and the kinematics of the components, as well as the mechanics (forces exerted), supplement the biochemical pathways in an essential manner^36-38^. For instance, it is now established that tension on the kinetochore, which attaches the chromosomes to the spindle apparatus, is critical to the spindle assembly checkpoint and to correct the erroneous attachments^39,40^. However, in human cells, such studies are still tour-de-force experiments. Furthermore, they require to synchronise the cells as the metaphase-to-cell-cycle duration ratio is about 5 %^41^. Model organisms, such as yeast, nematode, or drosophila, were instrumental in investigating the fundamentals of mitosis, as this ratio is much more favourable. The study of cellular division in mammalian cells often cannot be achieved by so-called physiological synchronisation (cell sorting, e.g.) or mechanical means like centrifugation. The classic workaround lies in synchronising cells by stopping their cell-cycle progression using drugs, often therapeutic molecules; washing it out lets the cells progress again, all at the same stage^42^. It appears sensible to block the cell cycle far from mitosis to decrease the impact of this treatment on mitosis mechanisms. Although more attractive, such strategies raised questions: while no major mitotic defect was observed, the progress of techniques to investigate dynamics and mechanics enabled us to uncover putative, more subtle perturbations^23^. For the sake of completeness, it is noteworthy that one can acquire a large number of asynchronous cells over a long time and select the mitotic cells a posteriori during analysis; beyond the waste of resources, such a strategy cannot be applied to study dynamics and mechanics as it requires imaging at several frames per second, which would cause photo-toxicity and photo-bleaching if applied over hours. Our Roboscope, by its autonomous ability to grab images of mitotic cells, is more parsimonious, preserving the sample and saving a lot of time and storage space while keeping the sample in physiological conditions.

More specifically, we wondered whether synchronisation may impact the mitotic spindle. In this respect, double thymidine block (DTB) synchronisation is attractive as it blocks the cells in the S phase, a distant phase from mitosis. Furthermore, this protocol is simple and affordable. However, whether such a protocol induces subtle defects is still an open issue^23^. Thus, we plated live cells, either synchronised or not, in similar conditions (Fig 3, Suppl Video 1). We then autonomously grabbed circa 30 metaphasic cells over 8 hours (metaphase probability above 0.9) without requiring the experimenter to intervene in each case.

**Figure 3:**
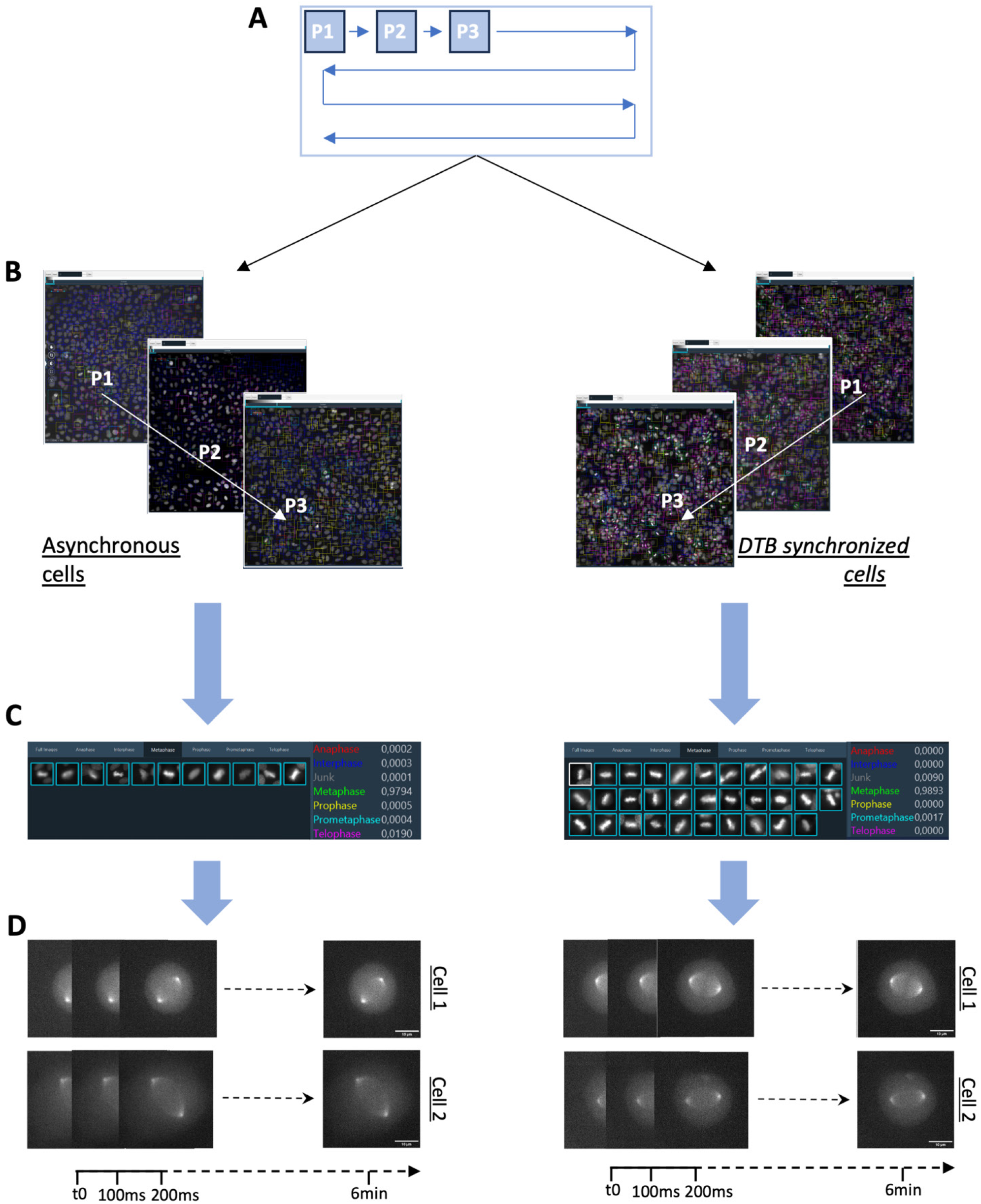
Metaphase-driven acquisition of spindles. (**A**) Plate diagram of acquisition. Sequence #1 screens every position registered at every time point and feeds images into the Roboscope module to detect metaphasic cells. (**B**) Typical images where some metaphasic cells were found in asynchronous (left) and mitosis-DTB-synchronized cells (right). (**C**) The *application* submodule then lists the candidates and their positions. (**D**) When the detected cell has a high enough probability of being in metaphase (threshold set when setting up the experiment), sequence #2 is started as exemplified. Such a sequence is run for each hit.

We tracked the images of the cells acquired at 10 frames per second during 205 seconds of metaphase finishing not later than 35 seconds before anaphase onset (set as the fast elongation of the spindle) using a modified version of our tracking algorithm successfully developed for the nematode^43,44^ (Fig 4A). We did not measure any significant difference in the average spindle length during that period, 12.9 ± 1.5 µm (mean ± SD, n = 7) for non-synchronised cells compared to 11.9 ± 0.8 µm (n= 7) for DTB synchronised ones (*p* = 0.15). This result was expected since such a protocol was used many times without observing major effects. However, the variation of length over time reflects the dynamicity of the spindle, for instance, the poleward flux of the microtubules^45^. We thus questioned the stability of spindle length. We band-pass filtered the time-traces between 0.015 Hz and 0.5 Hz using the robust local regression algorithm (rloess) to remove mean, residual drift and high frequency noise and computed the filtered variance similar to the filtered standard deviation^44^. We observed a net increase of the variance of spindle length upon DTB synchronisation (Fig 4D) while the variance of spindle transverse position remains grossly stable (Suppl Fig S8G). We suggest an effect of DTB synchronisation specific to the spindle length, which may reflect a weaker mechanical coupling of the two spindle poles.

**Figure 4:**
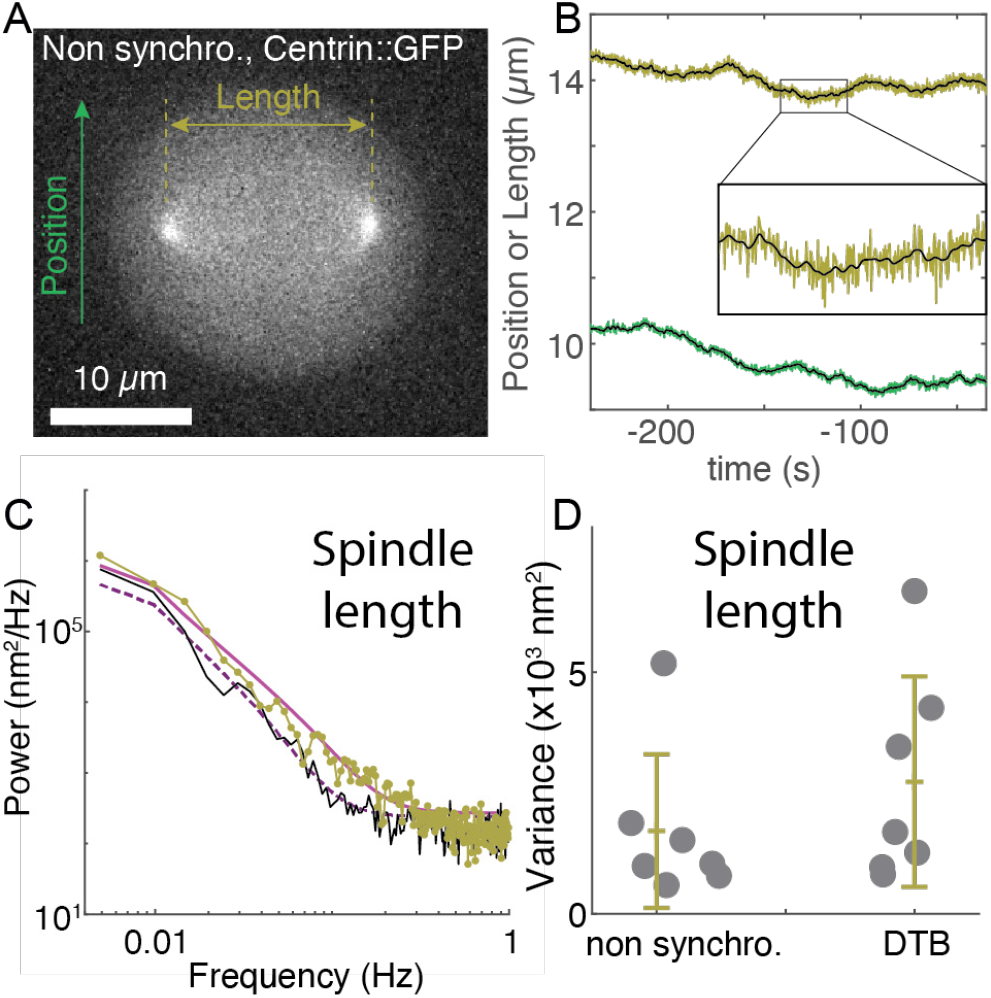
Spindle mechanics of U2OS cells synchronised or not with Double Thymidine Block (DTB) imaged with the Roboscope during metaphase. **(A)** Exemplar non-synchronised cell used for the subsequent analysis (the same cell as Suppl Video 1). The green arrow depicts the transverse axis for measuring the spindle centre position, and the khaki arrow schematises the spindle length measurement. (**B**) Spindle position along the transverse axis (green, arbitrary origin) and spindle length (khaki) measures considering a 205 s interval during metaphase. The overlaid black line is a 0.5 Hz low pass-filtered trace corresponding to the frequency range used when fitting the fluctuation spectra. Inset: magnified section of the spindle length curve. (**C**) Analysis of the fluctuations of the spindle length. Average power spectra from *n* = 7 non-synchronised (black line) and DTB-treated cells (khaki line). The magenta line are least-squares fits to the Kelvin-Voigt with inertia data-model for non-synchronised (dashed line) and DTB-treated cells (plain line). See corresponding parameters in Fig S8BD legend. (**D**) Variance along time of the spindle length for the same cells as for fluctuation analysis, band-pass filtered between 0.015 Hz and 0.5 Hz. Dots are the values for individual cells. Bars report the mean ± the standard deviation.

To further investigate this putative alteration, we measured the fluctuation of spindle length over time from the above tracks, and we computed the power spectral density (PSD, see Suppl Info for details) to recapitulate the mechanics of the spindle. We then fitted it with a Kelvin-Voigt model with inertia equivalent to second-order Lorentzian (Suppl Fig S8A-D)^44^. This model was convolved with Hann-window transform to account for the windowing^46^. Comparing averaged experimental PSD or their fits showed an up-shift of the spectra upon synchronisation consistent with a less rigid coupling between the spindle poles (Fig 4C). Importantly, we can exclude a batch or global effect as the spectra of the spindle transverse position were similar in both conditions (Suppl Fig S8E). We concluded that a DTB is likely to affect spindle mechanics. Using the Roboscope was essential to monitor the spindle dynamics without synchronisation and thus to reach this conclusion.

## Discussion

We report on the Roboscope, a new smart system enabling autonomous acquisition in fluorescence microscopy. The advantages of our approach are fourfold:

**Generic** through: (i) building up on our previous optimal driving, which controls a broad range of camera-based microscope setups and all their attached devices^16^; (ii) having a modular software architecture, in particular, enabling to change the image processing and the decision algorithm (two modules) to comply with a broad range of experiments, including beyond fluorescence microscopy; (iii) embedding a novel neural network, called sGAN, which could be easily adapted to a broad range of biological questions (El Habouz et al., in preparation). Such versatility makes the Roboscope highly suitable to be deployed in facilities.

**User-friendly** using a deep-learning analysis that can be adapted to a broad range of problems without computer coding but by transfer-learning or fine-tuning. The original design of our solution further reduced the number of cell vignettes required to less than 100 per class, and accuracy can be further improved by providing supplemental non-labelled images. Finally, our approach prevents biases introduced by EOI selection by the experimenter^47,48^. **Fast** through optimised microscope driving and sequence interruption, dedicated modules running in parallel, and a 20-layer-only neural network to classify images embedded on a specialised device. By doing so, we can capture transient dynamics and rare events that are proven essential to understanding the cell mechanisms^49-51^. The Roboscope, beyond efficiently coping with the mismatch between investigated events’ duration and their period of appearance^13^, will also fasten high content screening by focusing only on interesting EOI.

**Parsimonious**, by restricting illumination of the dyes of interest to EOI and fetching only qualified events to the computer. It prevents photo-damage by better managing the photon budget^52^. It also clearly saves the user time by avoiding extensive acquisition supervision and limiting the image storage requirements. In a broader take, this innovative system can be adapted to any biological issue as long as a small training set ensures the domain adaptation of the neural network or a fast-enough alternative network is provided.

As a proof-of-concept, we monitored spindle micro-movements with high frame rate acquisition during metaphase using the Roboscope. We found that, compared to non-treated cells, DTB-treated cells show altered mechanics despite having the same average spindle length. Detailed interpretation is out of the scope of this paper. However, because the strongest effect is a decrease of inertia in the model, it may be interpreted as higher dynamics of the component of the spindle in treated cells, for instance, faster growth/shrinking of microtubules, higher catastrophe/rescue rates, or faster binding/unbinding of molecular motors or microtubule cross-linkers. This would be consistent with putative DNA damages, or incorrect expression of some cell cycle proteins reported previously^53,54^. In the needed extra investigation, the Roboscope will be highly instrumental in easing the acquisition of non-synchronized metaphasic cells.

We showcased the Roboscope using one fluorescence channel for event detection and a second one to investigate spindle mechanics. While in the reported example, automation only changed acquisition conditions, its ability to retrieve cell position and feed it into the devices-driving system paves the way to photo-perturbative experiments. Beyond making tour-de-force experiments unbiased and no longer requiring tight supervision in fundamental research, it enables us to envision porting such experiments to High-Content Screening. It would greatly enlarge the way to get phenotypes, an essential aspect of screening, especially with the advent of machine learning analysis^55^. Conversely, the ability to conveniently generate large data sets is essential to the success of data-centred approaches^56^. Beyond research, autonomous microscopy also holds important promises for personalised medicine by enabling easy and standard examination of patient explants to choose the most appropriate treatment^57,58^.

## Online Methods

### Cell culture

Cells were cultured in Dulbecco modified Eagle medium (DMEM, Gibco, Life Technologies) supplemented with 10 % fetal calf serum (FCS) (Gibco, Life Technologies) and 1 % penicillin/streptomycin (100 U/ml, Gibco, Life Technologies) in a humidified atmosphere with 5 % C0_2_ at 37 °C. For synchronisation in mitosis, cells were treated with 2.5mM thymidine (Sigma-Aldrich) for 16h, released for 8h and then treated again with thymidine for 16h. After two washes with phosphate-buffered saline (PBS), cells were released for 10h according to the protocol described in ^59^.

For mitosis classification training sets, Hela cells (ECACC) were chosen. For spindle micro-movements analysis, U2OS cells (gift from G. Timinszky) were also transfected with eGFP-Centrin2^27^ (Nigg UK185 – Addgene) at a concentration of 1µg/µL with Xtreme-Gene HP (DNA Transfection reagent ratio 3:1, Merk) during 48 hours. Before image acquisition, DNA was stained with 0,1μg/ml Hoechst 33342 (H3570 - Thermo Fisher Scientific) for 1h at 37 °C.

### Image acquisition for mitosis classification

The training set was acquired on both Zeiss and Leica microscopes, while the reported application was done on the Leica only. Twenty thousand cells were distributed per well in glass-bottomed Imaging Chambers (Mo Bi Tec®) and cultured as previously explained. Each well was followed by time-lapse fluorescent microscopy in imaging media (L15 5 % FCS 1 % P/S). Acquisition lasted at least 6 hours (1 acquisition/10 minutes) inside a heating chamber. For both systems, resolution was 3.08 pixel/µm.

(<u>Zeiss microscope</u>) Images were acquired with an inverted widefield Axio Observer microscope fitted with a 0.8 NA 20X objective, a CoolSNAP CCD camera (Roper Scientific), a DAPI filter, and a 100 ms exposure time.

(<u>Leica microscope</u>) Images were acquired with an inverted widefield Leica DMi8 microscope fitted with a 0.4 NA 20X objective and a sCMOS camera OrcaFlash 4.0 V3 (Hamamatsu) with 100 ms exposure using a DAPI filter for screening.

### The Roboscope system

The Roboscope system, as described earlier, combines a Leica DMi8 microscope equipped with two dry objectives: 20x NA 0.4 and 63x NA 0.7, an ITK Hydra TT stage, an Nvidia AGX Xavier graphic adapter image processing, the Inscoper Imaging Solution and a middle range workstation. The microscope stage was inside a temperature control system (Life Imaging Services). Illumination light comes from a halogen lamp Leica EL6000 guided by a liquid fibre to the microscope. Two filter sets were used: one to image Hoechst (EX: 350/50, BS: 400, EM: 460/50; DAPI dichroic filter cube) and the other to image eGFP (Ex 480/40, BS: 505, EM: 527/30; GFP dichroic filter cube). Imaging was formed by an sCMOS camera OrcaFlash 4.0 V3 from Hamamatsu.

### Training of the deep network

The training was performed on two workstations: the first one had an Nvidia Tesla V100 (TensorFlow v 2.1) GPU for acceleration and was used for initial training; the second workstation had a GeForce RTX 2080 Ti (TensorFlow v 2.9) and was used for domain adaptation (Suppl Info).

### Centrosome tracking and analysis

The tracking of labelled centrosomes and analysis of trajectories were performed by custom tracking software^43,44^ and developed using Matlab (The MathWorks). Tracking of 20ºC methanol-fixed -tubulin-labelled C. *elegans* zygote indicated accuracy to 10 nm. In human cells, the high-frequency noise plateau in the power density spectrum indicated 148 nm2/Hz in the control condition and 158 nm2/Hz for DTB-treated cells, consistent with similar accuracy as observed in worm zygotes. Cell orientation and anaphase onset timing were achieved manually. The results were averaged over all of the replicas for each condition.

## Data availability

A GitHub repository (https://github.com/JacquesPecreaux/Roboscope_paper_Bonnet_et_all_24) archived on zenodo (http://doi.org/10.5281/zenodo.13832258 and http://doi.org/10.5281/zenodo.13832264) offers the annotation tools and dataset generated in the framework of this project.

## Acknowledgements

U2OS cells were kindly given by G. Timinszky, Munich, Germany. This project was supported by Région Bretagne (AAP PME 2018-2019 – Roboscope) and Agence Nationale de la Recherche (PRCE project SAMIC ANR-19-CE45-0011). Some microscopy imaging was performed at the Microscopy Rennes Imaging Center, UMS 3480 CNRS/US 18 INSERM/University of Rennes. We acknowledge France-BioImaging infrastructure supported by the French National Research Agency (ANR-10-INBS-04).

## Author contributions

Conceptualisation: OB, JP, MT; Methodology: JB, YEH, CM, OB, JP, MT; Software: YEH, CM, BG, BM; Validation: OB, JP, MT; Formal analysis: YEH, CM, JP; Investigation: JB, LR, YEH, CM, MG; Resources: JP, MT. Data Curation: JB, LR, CD, CM, JP, MT; Writing - Review & Editing: JB, JP, MT; Visualization: JB, JP; Supervision: OB, JP, MT; Project administration: OB, JP, MT; Funding acquisition: OB, JP, MT.

## Conflict of interest

CM, CD, and BG are employed by Inscoper, SAS. OB is the company’s chief technical officer. The Roul et al. 2014 patent on optimal driving is licensed exclusively to Inscoper, SAS. This company also co-owns with CNRS and the University of Rennes the Balluet et al. 2020 patent used in the reported system. JP and MT are the company’s scientific advisors. The funders had no role in study design, data collection and analysis, decision to publish, or manuscript preparation.

## Notes

http://doi.org/10.5281/zenodo.13832258

http://doi.org/10.5281/zenodo.13832264

