## Supplemental text, figures and tables for "The Roboscope: Smart and Fast Microscopy for Generic Event-Driven Acquisition"

The roboscope, as an autonomous microscope, needs to classify grabbed images on the fly to decide about the next acquisition steps. Thus, the algorithm must be fast enough to support catching transient and dynamical events, and in doing so, it benefited from embedding on a dedicated ARM-device GPU-accelerated. Our algorithm must also be generalisable to a broad range of labelling, imaging modalities, and biological questions without coding. This led us to use a Convolutional Neural Network (CNN) to detect the Events or Objects of interest (EOI). The limitation of such an approach is often the amount of training data (Greener et al., 2022). To keep the roboscope practicable, especially in microscopy facilities, we further required that it can be adapted to additional biological questions in a user-friendly fashion. It was achieved with a reduced training set by transfer learning or fine-tuning. In particular, we designed a semi-supervised Convolutional Neural Network (CNN) published at [El Habouz et al. in prep] inspired by (Odena, 2016) and termed sGAN. It could be trained by combining annotated and non-annotated images. On the grounds of this innovative network for autonomous microscopy, we detail the preprocessing of the images and the network training here. Finally, to showcase the roboscope, we characterised the mechanics of the mitotic spindle and compared synchronised and non-treated cells. We here further explain this approach.

#### 1 The sGAN is a generic and fast image classifier that fosters smart microscopy.

##### 1.1 Preprocessing of images:

###### 1.1.1 Image normalisation.

Both cameras used acquired images in size 2048x2048 pixels, 16 bits (Methods). These images contain 200 to 600 cells. The samples' diversity, the variability of labelling (due to transient fluorescence construct transfection, e.g.) or uneven illumination in the microscope field called for an image normalisation (Ioffe and Szegedy, 2015). We obtained the best results by adjusting the settings per microscope setup. In further detail, for images from the Zeiss microscope (Methods), we saturated the 0.5% darkest and brightest pixels by adjusting brightness and contrast on the entire image (before segmentation). For pictures from the Leica microscope (roboscope), we saturated no pixel, instead rescaling the image linearly so that the lower and higher intensities correspond to the minimum and maximum of the 16-bit encoded image. It is equivalent to setting 0% of pixels saturated in the algorithm above.

##### 1.1.2 Segmentation to produce the cell vignettes

Because the classifier cannot localise objects but classify single-cell images, we segmented the image to make sub-frames, called vignettes, each containing an individual cell. Although an extra deep-learning-based algorithm could achieve this task, we set out to use an old-style algorithm for the sake of simplicity of the prototype. Importantly, for isolated cells, it turns out that this algorithm is generic enough to perform by simply adjusting the hyperparameters once and for all to adapt to various microscopes and cell types, provided nuclei are labelled and cells are not too densely packed. The core of this algorithm is adaptive thresholding and a watershed to separate contacting cells (Gonzalez and Woods, 2002; Vincent and Soille, 1991).

We implemented it using the OpenCV package (www.opencv.org) as follows:

1- The input image was slightly smoothed using *GaussianBlur* with the following parameters: *gaussian_kernel_size* = 3, *gaussian_filter_sigma* = 4.
2- The image was then transformed into a binary image using the *adaptiveThreshold* parameterised with *threshold_block_size* = 85 and *threshold_const* = -2,
3- And denoised with morphologyEx.

At this stage, the objects were roughly visible and could be segmented.

4- The *distanceTransform* method (derived from watershed) followed by *adaptiveThreshold* determined highlighted the pixels belonging to the object.
5- We computed its boundary by subtracting the images from (3) and (4).
6- We finally applied the watershed algorithm to separate objects correctly.

The values reported here are those used for the Leica/Roboscope system during reported experiments. To generate a segmentation ground truth, the Zeiss and Leica training datasets were segmented manually (and classified) by experts, not with this algorithm.

#### 1.2 Semi-supervised GAN:

##### 1.2.1 Architecture

We designed the sGAN as a combination of a Generative Adversarial Network (GAN) (Goodfellow et al., 2014), an unsupervised method, and a supervised CNN classifier (Odena, 2016). The supervised classifier part of our SGAN shared the same weights with GAN's discriminator, except for the last layer (Fig S1). In doing so, we improved the training by supplementing the labelled training set with non-labelled images, decreasing experts' burden due to annotation. Augmented by the synthetic images from the GAN generator network, it made the training threefold more data thrifty. Such a training strategy also increases the generalisability of our network. Interestingly, the discriminator and supervised classifier were only 20-layer depth (Table S1), which ensured fast classification when embedding into the roboscope and reduced the training set's size.

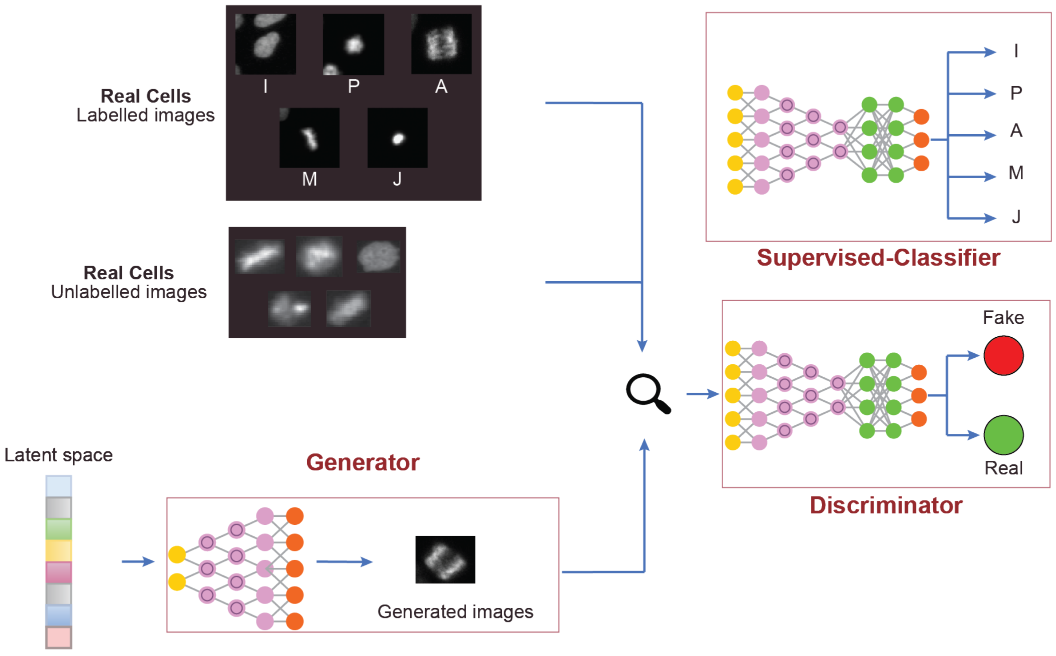


**Figure S1:** Architecture of the proposed deep learning network, termed sGAN. It features three major components: (**a**) The supervised classifier, which takes an image as input and returns its class. The last layer of this classifier is trained on labelled images while the other layers parameters are copied from the discriminator; (**b**) The generator, which receives a Gaussian vector of size 100 and generates an image; (**c**) The discriminator, taking a generated or real image and returning a True/False answer whether this image is real. The images used by this latter are not labelled.

| Layer Type | Output Shape | Number of parameters |
|---|---|---|
| Conv2D | (72,72,64) | 1792 |
| Conv2D | (72,72,64) | 36928 |
| MaxPooling2D | (36,36,64) | 0 |
| Conv2D | (36,36,64) | 73856 |
| Conv2D | (36,36,64) | 147584 |
| MaxPooling2D | (18,18,128) | 0 |
| conv2D | (18,18,256) | 295168 |
| conv2D | (18,18,256) | 590080 |
| conv2D | (18,18,256) | 590080 |
| MaxPooling2D | (9,9,256) | 0 |
| Conv2D | (9,9,512) | 1180160 |
| Conv2D | (9,9,512) | 2359808 |
| Conv2D | (9,9,512) | 2359808 |
| MaxPooling2D | (4,4,512) | 0 |
| Flatten | (8192) | 0 |
| Dropout | (8192) | 0 |
| Dense | (1000) | 18433000 |
| Dropout | (1000) | 0 |
| Dense | (800) | 800800 |
| Dense | (7) | 5607 |

| | |
|---|---|
| **Total parameters** | **16634671** |
| **Total trainable parameters** | **16634671** |
| **Total non-trainable parameters** | **0** |

**Table S1:** Architecture of our supervised GAN (supervised classifier of Fig S1).

##### 1.2.2 Implementation

At training time, we executed the network on a Python implementation of TensorFlow version 2.1 running on a high-range workstation equipped with an NVidia Tesla V100 GPU. Subsequent transfer learning or fine-tuning was done on a second workstation with a GeForce RTX 2080 Ti and TensorFlow v 2.9. At run time, we used a Python implementation of TensorRT and converted the network appropriately. We considered only the supervised-classifier part on the sGAN and discarded the others. This network was loaded on a Nvidia AGX Xavier device with the weights from the training described below.

#### 1.3 Datasets and annotation

##### 1.3.1 Mitocheck

We used the database termed *mitocheck* (Neumann et al., 2010) for full training, prepared similarly to (Balluet et al., 2022). This dataset comprises wide-field fluorescence time-lapses of Hela Kyoto cells, expressing chromatin GFP marker, acquired with a 10x dry objective on Olympus ScanR. Several mitotic phases and defect phenotypes were observed. We only considered a subset of 5 classes: interphase, prometaphase, metaphase, anaphase, and apoptosis. Note that junk images in our experiments usually end up classified as apoptosis. We also equilibrated the dataset (Fig S2). Doing so, it finally contained 685 images cropped to 72x72 pixels.

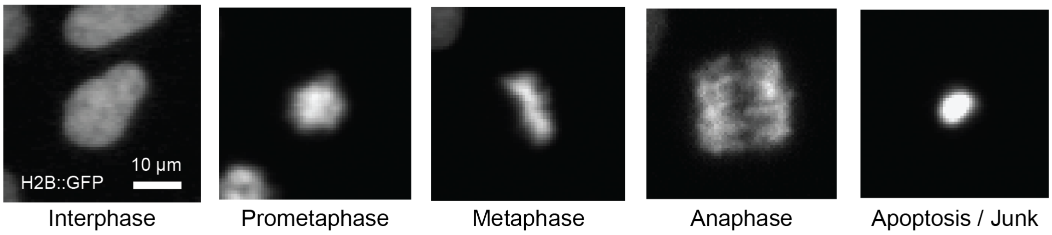


| Class | Size | Data-split | | Marker | Cell line | Details | Source |
|---|---|---|---|---|---|---|---|
| | | Train | Test | | | | |
| Interphase<br>Prometaphase<br>Metaphase<br>Anaphase<br>Apoptosis / Junk | 685 | Labelled: 400<br>Unlabelled: 185 | 100 | Histone GFP tagged | HeLa (human) | Wide-field fluorescence time-lapse microscopy with 10x dry objective on Olympus ScanR. | Neumann et al. 2010 |

**Figure S2**: Mitocheck dataset used for initial training of the proposed algorithm sGAN. (top): Exemplar images of each class. (bottom): Statistics and information on the dataset.

##### 1.3.2 Zeiss and Leica/Roboscope datasets

Barely using the initially trained network resulted in poor classification [El Habouz, *in prep*]. We set out to perform domain adaptation with images acquired directly on the roboscope (Leica) system. We supplemented these images with some acquired using a Zeiss microscope to get further genericity. Notably, typical image histograms differed between these two systems. We prepared Hela Kyoto cells on both systems, and DNA was labelled with Hoechst (Methods). We imaged these cells with 20x dry objectives. To get enough transient classes and equilibrate the dataset, the cells were synchronised by a double thymidine block (Methods). The produced dataset contained 2126 cell vignettes of size 72x72 manually classified into seven classes: interphase, prophase, prometaphase, metaphase, anaphase (early and late), telophase, and apoptosis. Notably, circa 80%

of the cells in both datasets were classified as interphase despite synchronisation. To keep the dataset equilibrated, not all interphase cells or cells from other abundant classes were annotated. To this end, the script *balanceDataset.py* was instrumental (Data availability).

The **Leica dataset** features ten time-lapse experiments featuring 19 time-points imaging double thymidine block synchronised cells. It was supplemented with eight experiments featuring seven time-points imaging unsynchronised cells. Two experts annotated all the experiments, and only vignettes identically classified by both were kept. This latter procedure was automatised by the script *mergeAnnotations.py* (Data availability; Methods). These images were 16-bit encoded 2048x2048 pixels sized. We finally obtained 115 vignettes per class. (Fig S3, Data availability, subfolder *balanced_Leica_Dataset*).

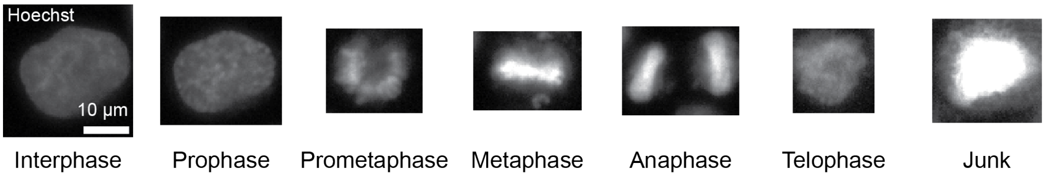


| Class | Size | Data-split | | Marker | Cell line | Details | Source |
|---|---|---|---|---|---|---|---|
| | | Train | Test | | | | |
| Interphase<br>Prophase<br>Prometaphase<br>Metaphase<br>Anaphase<br>Telophase<br>Junk | 1321 | Labelled: 985<br>Unlabelled: 0 | 336 | DNA with Hoechst 33342 | HeLa (human) | Wide-field fluorescence time-lapse microscopy with 20x dry objective on Zeiss Axis Observer. | This study |

**Figure S3:** Zeiss dataset used for full training. (top): Exemplar images of each class. (bottom): Statistics and information on the dataset (Methods).

The **Zeiss dataset** features a single time-lapse experiment featuring 31 time-points imaging double thymidine block synchronised cells. This 2048x2048 pixels 16-bit image stack was broken into four tiles 1024x1024 pixels-sized, annotated separately and used as subsets for cross-validation. One expert annotated half of the experiments, and the other half was annotated by the other. We finally obtained 192 vignettes per class, except for the junk class with 169 (Fig S4, Data availability, subfolder *balanced_Zeiss_Dataset*; Methods).

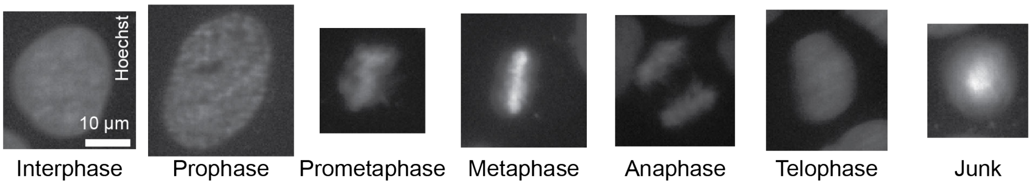


| Class | Size | Data-split | | Marker | Cell line | Details | Source |
|---|---|---|---|---|---|---|---|
| | | Train | Test | | | | |
| Interphase<br>Prophase<br>Prometaphase<br>Metaphase<br>Anaphase<br>Telophase<br>Junk | 805 | Labelled: 525<br>Unlabelled: 0 | 280 | DNA with Hoechst 33342 | HeLa (human) | Wide-field fluorescence time-lapse microscopy with 20x dry objective on Leica DMi8. | This study |

**Figure S4:** Roboscope (Leica) dataset used for fine-tuning. (top): Exemplar images of each class. (bottom): Statistics and information on the dataset (Methods).

##### 1.3.3 Annotation

The images were annotated using the software labelImg (Tzutalin, 2018). The software allowed us to annotate entire images, localise cells, and choose their class. Annotations were saved in .xml files. To feed into the sGAN training, we extracted cell vignettes using a homemade labelImg modification, especially the function "create slice" (Data availability).

#### 1.4 Training

##### 1.4.1 Training parameters and data augmentation

The parameters were manually optimised to achieve full training using the Zeiss dataset training and fine-tuning using the Leica dataset. They read:

- Batch size: 4
- Loss function: KLDivergence
- Optimised: SGD, learning rate: 0.0001
- Data augmentations :
  - Rotation range: 20
  - Width shift range, height shift range: 0,1
  - Brightness range : [0.9,1]
  - Zoom range : [1.0,1.3]

Fill mode: nearest

The validation dataset's loss function is monitored during training to perform an early stop when loss stops decreasing.

Note that training using the Mitocheck dataset departed from these settings in not using brightness and zoom data augmentation and using *categorical_crossentropy* as a loss function.

##### 1.4.2 Training strategy

**Initial training from scratch**

We fully trained the sGAN described above using randomly initialised weights and the Mitocheck dataset. We split the dataset into 400 labelled images for training, 185 presented without any label to the algorithm, and 100 images for testing. We obtained an accuracy of 86% (Fig S5). The training lasted 6 hours using the NVidia V100 GPU.

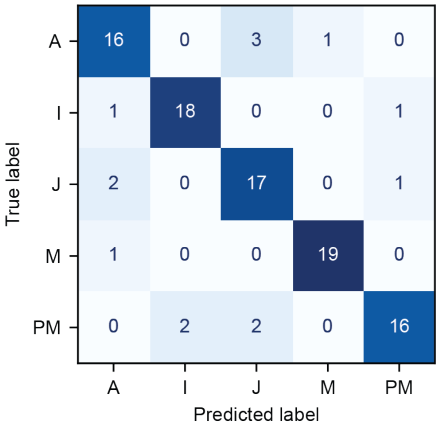


| Class | Precision | Recall | F1-Score |
|---|---|---|---|
| Anaphase | 80 % | 80 % | 80 % |
| Interphase | 90 % | 90 % | 90 % |
| Junk / Apoptosis | 77 % | 85 % | 81 % |
| Metaphase | 95 % | 95 % | 95 % |
| ProMetaphase | 89 % | 80 % | 84 % |

Accurary: 86 %

**Figure S5**: Full training of our SGAN from randomly initialised weight using the mitocheck dataset. (left) The confusion matrix on the 100 test images, with true labels in rows and predicted ones in columns. (right) Statistics of trained classifier, per class.

##### Full retraining from the initialised network using the Zeiss dataset.

We retrained with a second dataset to increase the diversity of training images. Considering the four tiles of size 1024x1024 pixels (see §1.3.2), we used 3 to extract 144 vignettes per class for training (121 for *junk*) and one as a test set (48 vignettes per class). The weights of the sGAN were inherited from initial training with Mitocheck. We performed 4-fold cross-validation; we obtained an accuracy of 83.1 ± 1.1% (Fig S6). Except for some confusion between Interphase and Telophase, the results appeared sufficiently accurate for the targeted applications. Using the NVidia GeForce RTX 2080Ti GPU, the training lasted 29 minutes.

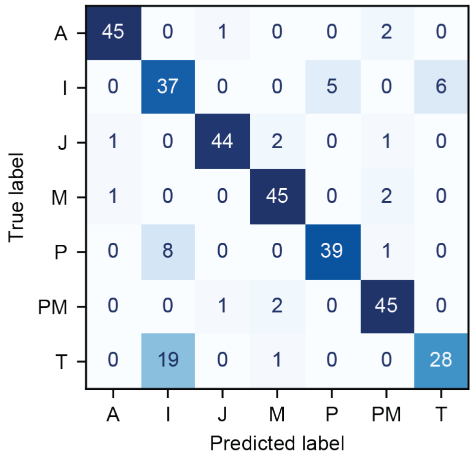


| Class | Precision | Recall | F1-Score |
|---|---|---|---|
| Anaphase | 96 % | 94 % | 95 % |
| Interphase | 58 % | 77 % | 66 % |
| Junk / Apoptosis | 96 % | 92 % | 94 % |
| Metaphase | 90 % | 94 % | 92 % |
| Prophase | 89 % | 81 % | 85 % |
| ProMetaphase | 88 % | 94 % | 91 % |
| Telophase | 82 % | 58 % | 68 % |

Accurary: 84 %

**Figure S6**: Full training of our SGAN from mitocheck-trained weights using the Zeiss dataset and 4-fold cross-validation. (left) Typical confusion matrix on 336 test images, with true labels in rows and predicted ones in columns. (right) Statistics of trained classifier, per class.

##### Domain adaptation: fine-tuning using the Leica (Roboscope) dataset.

We finally fine-tuned images taken on the Roboscope (Leica) system to optimise the detection. We divided the dataset by keeping either 65% (75 vignettes per class) or 83% (95 vignettes per class) for training and the remaining for testing (38 or 20 vignettes per class, respectively). We performed a fine-tuning using 3-fold cross-validation and obtained an accuracy of 75.2 ± 2.7% (Fig S7). Surprisingly, using 6-fold cross-validation to increase the number of images used in training, accuracy read 74.1 ± 3.8%, not better than before. A full retraining (from the initialised network) does not improve this value. It is noteworthy that for instrumental reasons, at the time of training the roboscope for the reported experiments, the image quality was lower than reported here, although sufficient to perform experiments. We improved the instrument in the meantime and set out to report on its current status. Using the NVidia GeForce RTX 2080Ti GPU, the training lasted 36 minutes.

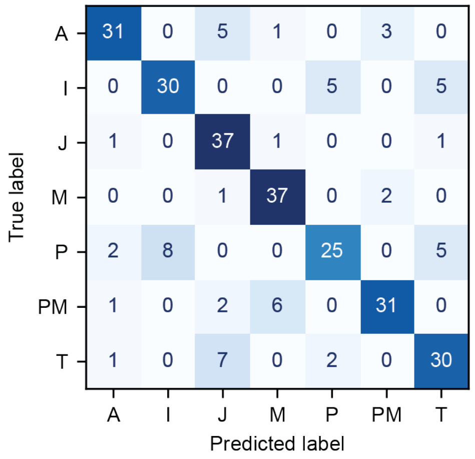


| Class | Precision | Recall | F1-Score |
|---|---|---|---|
| Anaphase | 86 % | 78 % | 82 % |
| Interphase | 79 % | 75 % | 77 % |
| Junk / Apoptosis | 71 % | 93 % | 80 % |
| Metaphase | 82 % | 93 % | 87 % |
| Prophase | 78 % | 63 % | 69 % |
| ProMetaphase | 86 % | 78 % | 82 % |
| Telophase | 73 % | 75 % | 74 % |

Accurary: 79 %

**Figure S7**: Fine-tuning of our SGAN from mitocheck- then Zeiss-trained weights using the Leica/Roboscope dataset and 3-fold cross-validation. a) Typical confusion matrix on 280 test images, with true labels in rows and predicted ones in columns. (b) Statistics of trained classifier, per class.

### 2 Spindle fluctuation analysis in living cells

#### 2.1 Power density spectrum as a mechanical fingerprint of the spindle

We set to compare the spindle length maintenance mechanism in the context of double-thymidine-block synchronised and non-synchronized cells. To do so, we investigated the fluctuation of spindle length over time based on our previously published method (Pecreaux et al., 2016). The obtained power spectral density (PSD) can be viewed as the variance-of-length-variations distribution over frequencies. In contrast to our previous publication, the spindle length changed over time; we cannot assume equal length at the beginning and the end of the considered interval. Since the discrete Fourier transform (DFT) used to compute the spectra assumes the periodicity of the signal, we applied a multiplicative window to safeguard against artefacts, as classically performed in signal processing (Mercat, 2016; Press, 2007). As a result, the obtained PSD was calculated from the DFT of spindle-length fluctuations convolved with the DFT of the window. We opted for the Hann window, which displays only three non-zero values in Fourier space, minimally altering the spectrum (Harris, 1978). We then repeated such a PSD computing over several cells to lower the error bars on the experimental spectrum by averaging and computing error bars (Fig S8A-D, black circles) (Berg-Sorensen and Flyvbjerg, 2004).

We observed an up-shift of the PSD of spindle length upon DTB synchronisation consistent with a less rigid coupling between the spindle poles (Fig 4C). To exclude a global effect, we repeated the analysis comparing the spindle transverse position and did not see a similar PSD shift (Fig S8E). We concluded that there was an effect specific to spindle length.

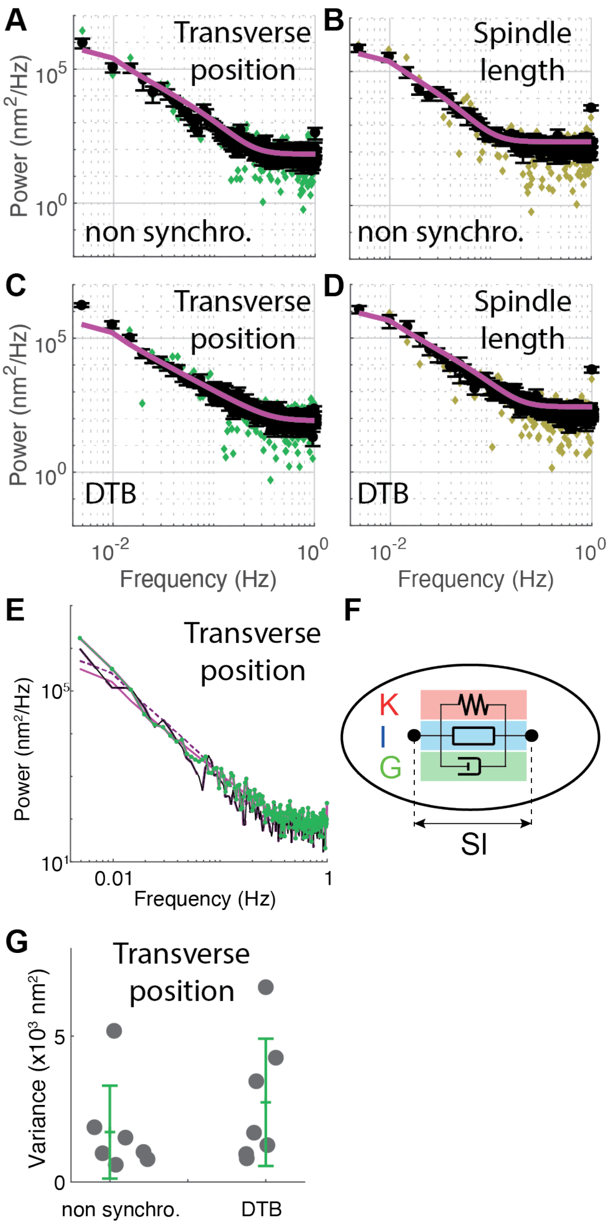


**Figure S8:** Analysis of the fluctuations of spindle transverse position and length of U2OS cells synchronised or not with Double Thymidine Block (DTB) imaged with the roboscope during metaphase. (**A-D)** Experimental and theoretical power spectra. (**A**) (Green diamonds) One-sided power spectral density of the transverse position of the spindle computed during the metaphase for a typical non-synchronised cell (the same as Fig. 4). (Black line) Average of power spectra from $N = 7$ cells. (Solid magenta line) Least-squares fit to the Kelvin-Voigt with inertia data-model with $K = 4.1 \pm 80$ µN/m, $G = 650 \pm 90$ µN.s/m and $I = 890 \pm 10$ µN.s²/m. (**B**) (Khahi diamonds) Corresponding one-sided power spectral density of the spindle length for the same cells, with fit to the Kelvin-Voigt-with-inertia data-model with $K = 8.9 \pm 39$ µN/m, $G = 660 \pm 80$ µN.s/m and $I = 2340 \pm 3$ µN.s²/m.

(**CD**) Similar plots for $N = 7$ DTB synchronised cells. Fit parameters read for transverse position $K = 0.59 \pm 831$ µN/m, $G = 740 \pm 75$ µN.s/m and $I = 300 \pm 31$ µN.s²/m and for spindle length $K = 7.1 \pm 27$ µN/m, $G = 560 \pm 50$ µN.s/m and $I = 850 \pm 49$ µN.s²/m. Error bars for all averaged spectra are standard errors. (**E**) Average power spectra from the same (Black line) non-synchronised and (khaki line) DTB-treated cells. (Magenta line) Least-squares fit to the Kelvin-Voigt with inertia data-model for (dashed line) non-synchronised and (plain line) DTB treated cell with parameters above. (**F**) Schematics depicting the Kelvin-Voigt-with-inertia discrete model used for fitting spindle length and position.

#### 2.2 Fitting the power density spectrum and estimating the variance of the fluctuations

To get some insight, we fitted the power spectra with a Kelvin-Voigt-with-inertia discrete model corresponding to a Hookean spring, a dashpot, and an inertial element in parallel (Fig S8F). The corresponding Langevin equation read:

$$I\ddot{y} + G\dot{y} + Ky = \eta$$

*I* was the inertia related to the dynamics of components (Pecreaux et al., 2016), *G* was the damping, and *K* was the stiffness of the spring. $\eta$ corresponded to a stochastic force as discussed in (Pecreaux et al., 2016). This generic model was chosen because it can recapitulate a variety of cytoskeleton and molecular motor-based dynamics (Mercat, 2016). Furthermore, this model appeared simple and fitted our data well. We named it a data model for these reasons. Finally, it is equivalent to the second-order Lorentzian used to model spindle centring in the nematode C. *elegans* (Pecreaux et al., 2016). Such a model is transformed into the Fourier space and convolved on the fly with the transform of the Hann window to perform the fit (Mercat, 2016). We validated that the model correctly recapitulated the data by computing the variance of the spindle position or length over the time range of interest on the one hand and, on the other hand, by numerically integrating the adjusted data model over the frequency range considered (Table S2). We expected both values to be close, on the ground of the Parseval-Plancherel identity. Estimating variance through the experimental approach, more direct but sensitive to drift, or through the data-model fit of the PSD, less direct but insensitive to drift, suggested that DTB may cause the spindle to fluctuate more.

| Time-Variance of the signal ($\times 10^3$ nm$^2$) Mean ± s.e. | Non synchronised (*N* = 7) | | DTB treated (*N* = 7) | |
|---|---|---|---|---|
| | Experimental | Data-model based | Experimental | Data-model based |
| Transverse position | 38.5 ± 13.9 | 43.3 ± 27.0 | 91.9 ± 64.3 | 62.7 ± 26.9 |
| Length | 23.5 ± 4.3 | 40.5 ± 25.4 | 56.1 ± 18.6 | 56.7 ± 25.3 |

**Table S2**: Variances estimated using direct space "classic" estimator and data-model based estimator through PSD fingerprinting. Values are reported for spindle length and transverse position, for cells synchronised using DTB or not.

---

**Supplemental Video 1:** Exemplar cell non-synchronised labelled using Centrin::GFP and used as examples in Fig 4, imaged at 10 frames per second. The movie is ten times accelerated.
